# Chemotherapy response prediction with diffuser elapser network

**DOI:** 10.1101/2020.10.14.339010

**Authors:** Batuhan Koyuncu, Ahmet Melek, Defne Yilmaz, Mert Tuzer, Mehmet Burcin Unlu

## Abstract

In solid tumors, elevated fluid pressure and inadequate blood perfusion resulting from unbalanced angiogenesis are the prominent reasons for the ineffective drug delivery inside tumors. To normalize the heterogeneous and tortuous tumor vessel structure, antiangiogenic treatment is an effective approach. Additionally, the combined therapy of antiangiogenic agents and chemotherapy drugs has shown promising effects on enhanced drug delivery. However, the need to find the appropriate scheduling and dosages of the combination therapy is one of the main problems in anticancer therapy. Our study aims to generate a realistic response to the treatment schedule, making it possible for future works to use these patient-specific responses to decide on the optimal starting time and dosages of cytotoxic drug treatment. Our dataset is based on our previous in-silico model with a framework for the tumor microenvironment, consisting of a tumor layer, vasculature network, interstitial fluid pressure, and drug diffusion maps. In this regard, the chemotherapy response prediction problem is discussed in the study, putting forth a proof-of-concept for deep learning models to capture the tumor growth and drug response behaviors simultaneously. The proposed model utilizes multiple convolutional neural network submodels to predict future tumor microenvironment maps considering the effects of ongoing treatment. Since the model has the task of predicting future tumor microenvironment maps, we use two image quality evaluation metrics, which are structural similarity and peak signal-to-noise ratio, to evaluate model performance. We track tumor cell density values of ground truth and predicted tumor microenvironments. The model predicts tumor microenvironment maps seven days ahead with the average structural similarity score of 0.973 and the average peak signal ratio of 35.41 in the test set. It also predicts tumor cell density at the end day of 7 with the mean absolute percentage error of 2.292 ± 1.820.

**Author summary:** The disorganized structure and leakiness of tumor vessels induce the inadequate blood supply and high fluid pressure within tumors. These features of the tumor microenvironment, identified as delivery barriers, lead to an insufficient amount of drugs to reach the interior parts of tumors. It is observed that the use of anti-vascular drugs makes the structure and function of the tumor vascular system more normal. Moreover, the combination of these drugs with cytotoxic agents provides favorable results with increased treatment response. But, it is also important to adjust the treatment schedule properly. In this regard, we build a deep learning model, designed to examine the tumor response with the ongoing treatment schedule. Our study suggests that deep learning models can be used to predict tumor growth and drug response in the scheduling of cytotoxic drugs.

## Introduction

Tumors need their blood supplies to grow beyond the size of 1-2 mm^3^ in diameter and meet the needs of oxygen and other nutrients. For this reason, tumors stimulate angiogenesis, a process in which tumors form the new blood vessels from pre-existing ones by secreting the various growth factors and, most importantly vascular endothelial growth factor (VEGF). The angiogenic switch between proangiogenic and antiangiogenic factors is activated for tumor progression and metastases [1]. Due to tumor-induced angiogenesis, the newly developed vessels have a leaky and disorganized structure accompanying a microenvironment, identified by hypoxia, acidosis, and increased fluid pressure [2]. As a result, this structurally and functionally abnormal tumor vascular network leads to the heterogeneous and inadequate drug distribution inside tumors.

The chaotic architecture, high vascular permeability of tumor vessels, and the lack of functional lymphatics lead to elevated interstitial fluid pressure (IFP), creating a barrier for the transport of therapeutic agents and nanoparticles [3,4]. IFP is uniform across the tumor nearly equal to microvascular pressure (MVP), but it drops precipitously at the tumor boundary. Therefore, the absence of a pressure gradient along vessels hampers the penetration of cytotoxic drugs to the interior parts of tumors transported by convection [3].

Normalization of tumor vasculature by using antiangiogenic agents is a widely used treatment modality in cancer therapy. Antiangiogenic agents maintain the balance between proangiogenic and antiangiogenic factors similar to healthy tissues. Furthermore, these agents normalize the structure and function of tumor vascular network transiently by inducing reduced vessel diameter and decreased vessel wall permeability. This process triggers tumor vessels to be more useful for the delivery of drugs as well as oxygen and nutrients to the targeted cancer cells [2,5]. Vascular normalization improves the convective transport of drug particles with a decrease in IFP, thereby inducing pressure gradients across vessel walls [1,6].

Antiangiogenic agents can act directly on the tumor vasculature [7]; also, several preclinical and clinical studies show that the application of antiangiogenic agents together with chemotherapy drugs provide beneficial results with increased therapeutic outcome [8–10]. The combination therapy can enhance the delivery of therapeutic agents to the interior parts of tumors [6]. Normalization is a transient process that means there is a time window for vessel normalization to occur. For this reason, chemotherapy drugs should be carefully administered within this window to benefit from the improved vessel conditions [11]. Moreover, the excessive application of antiangiogenic agents for more extended periods brings about vessel pruning, which decreases the outcome of combined therapies [12, 13]. Therefore, the appropriate timing and dosing of the agents should be carefully adjusted to improve the functionality of blood vessels as well as the delivery of anticancer drugs to tumor cells.

Mathematical models are commonly studied in cancer therapies to simulate the delivery of cytotoxic drugs and understand the relations between tumor microenvironment and drug delivery. Different approaches to the modeling of tumor vasculature and angiogenesis have been suggested to study antiangiogenic agents’ applications combined with chemotherapy drugs. By using discrete vasculature models, the delivery of chemotherapy drugs to tumors and the treatment response have been investigated extensively [14–17]. Also, in 2D and 3D blood flow models, the use of antiangiogenic agents and their effects on tumor response have been studied by Stephanou *et al.* [18]. Normalization is also simulated to investigate its effects on blood flow and the combination of antiangiogenic agents with cytotoxic drugs [13,19–21]. In addition to mathematical models, angiogenesis imaging is largely studied to examine tumor growth, characterization of tumor vasculature, and the response of the therapies. Employing various imaging modalities such as computer tomography (CT) and magnetic resonance imaging (MRI) gives an insight into the progress of tumor and vascular structure [22,23]. Additionally, photoacoustic imaging is exploited to monitor tumor angiogenesis due to its higher contrast and spatial resolution [24–26]. Besides, it is feasible to detect the early tumor growth and vascularization and observe the progression of antiangiogenic treatments [27, 28].

Early tumor response prediction, a measure of the effectiveness of treatment, is crucial in anticancer therapies to determine appropriate treatment schedules, apply the optimal drug dosages to patients, and increase patient survival. In this regard, to develop successful models for tumor response prediction, it is vital to monitor the distribution of cytotoxic drugs and to evaluate the tumor response to therapies in advance. Response prediction problems are often formulated by developing mathematical models benefiting from reaction-diffusion frameworks [29–31]. These models are capable of modeling tumor response to an extent, but they are generally deterministic models driven by a limited number of parameters that might constrain the model to comprise the inherent tumor growth patterns. However, several studies in deep learning outperform the traditional approaches in various tasks such as tissue classification, tumor segmentation, and tumor growth prediction [32–35]. A convolutional neural network (CNN) is developed by Urban *et al.* to classify the images of vascular networks taken before and after various antiangiogenic drug applications *in vitro* [36]. In another study led by Ha *et al.,* a CNN algorithm is used to determine the chemotherapy response prediction in patients with breast cancer. The breast MRI dataset is utilized as a baseline, and the treatment response before the application of chemotherapy drugs is predicted [37]. Positron emission tomography (PET) and CT images from different types of cancer are benefited in recent studies to develop CNN models that have the potential to predict the response of chemotherapy with a high sensitivity [38, 39].

Deep learning allows us to build computational models to learn representations of data with multiple levels of abstraction for the given task [40]. For instance, a feed-forward neural network is a function approximation that maps input data to output data. The function is formed by composing simpler non-linear functions where each function provides a new representation of input data [41]. After the model extracts the information out of the input data, it amplifies the essential features for a given task such as classification, segmentation, and regression. Convolutional neural networks, a neural network type, perform convolutions over the input by using the given number of filters, which are capable of learning the spatial and temporal dynamics in image data [40]. They are robust to overfitting due to the convolution operation properties, which reduces the full connection of the network. These models have applications in image [42] and video recognition [43], medical image analysis, and image processing [44]. By employing CNNs, models can be created for spatial and temporal forecasting problems in an end-to-end manner in the presence of sufficient data.

Tumor response prediction can be formulated as a spatio-temporal forecasting problem under the effects of interventions. Interventions can be described as drug injections. Therefore, the forecasting problem is equivalent to modeling the tumor microenvironment during the ongoing treatment. The tumor microenvironment can be described with tumor density, vasculature, interstitial fluid pressure (IFP), antiangiogenic treatment, chemotherapy maps, and drug dosages from the ongoing treatment. The model, denoted by *F*, takes tumor microenvironment maps *X_t_* and drug scalars *S_t_* as inputs at time *t* and predicts future tumor microenvironment maps *X_t+1_* at time *t* + 1. The predicted future tumor microenvironment maps allow us to investigate tumor growth and shrinkage patterns as well as drug diffusion maps, which are the key indicators of treatment efficiency.

In this study, we develop a deep learning model, which can capture the tumor response behavior conditioned by the ongoing treatment schedule. Our study suggests that deep learning models can be used to predict tumor growth and drug response in the scheduling of cytotoxic drugs. Since the required input data such as tumor density, IFP, vasculature, and drug maps is hard to collect from clinical patients for various reasons, we use the synthetic data from the mathematical model built in our previous paper [21]. Therefore, the proposed deep learning model *F* encapsulates the non-linear partial differential equations (PDEs) that govern the spatio-temporal dynamics of the tumor microenvironment. The model consists of multiple CNNs. Our motivation for using CNNs is based upon two key reasons; CNNs can extract coupled spatial features from multichannel inputs, and they can utilize the spatial features to predict future microenvironment maps. In the end, our model uses tumor microenvironment channels and drug scalars as inputs and predicts future tumor microenvironment maps which may assist to find appropriate scheduling and dosages of the combination therapy for the patients.

## Results

Our deep learning model aims to predict chemotherapy response by outputting future tumor microenvironment maps. The proposed model, Diffuser-Elapser Network (DENT), consists of seven submodels where each of them is a CNN. All submodels are trained separately as their tasks are different from each other. In the training, we perform cross validation among the unique patients. Therefore, we ensure that there is not any dependency between training and validation sets. Each submodel contains several convolutional layers. Model specifications and training procedures can be found in Methods section.

The DENT model takes input tensors describing the current tumor microenvironment state, which consists of five channels, namely tumor density, vasculature, IFP, antiangiogenic drug, and chemotherapy drug maps that are accompanied by chemotherapy and antiangiogenic drug dosages and predicts the future tumor microenvironment maps. The model can use its predictions as its next inputs. Since the model predicts future tumor microenvironment maps, we use two image quality evaluation metrics, which are peak signal-to-noise ratio (PSNR) and structural similarity (SSIM) [47]. PSNR represents a measure of the quality of prediction by calculating the ratio between the square of maximum fluctuation among pixels in the ground truth image and MSE between ground truth and predicted images. SSIM measures the perceptual difference between two images in terms of structural information in the images. In this method, ground truth images are considered as references that have the reference quality, and the quality of another image is measured by comparing it with the initial one. SSIM is mostly applied to improve or track the perceptual metrics based on the structural information; on the other hand, PSNR is based on estimating pixel-wise error. Both metrics are unitless quantities. The PSNR score approaches infinity as the MSE approaches zero. The range for an SSIM score is the interval of [−1,1]. A higher score denotes a better prediction quality in both metrics.

In the study, we first investigate the performance of the model on predicting future tumor microenvironment maps. To give an insight, we pick a synthetically generated case that contains a patient’s tumor microenvironment maps during treatment. The tumor microenvironment maps consist of five channels correspond to tumor cell density, vasculature, IFP, antiangiogenic drug diffusion, and chemotherapy drug diffusion maps. The input, ground truth, and predicted tumor microenvironment maps during the treatment are presented in Fig 1. We prefer eight days-long cases so that we can compare our predictions with the previous simulation findings [21], which is limited to simulating 8 days after the initiation of therapy. The patient receives treatment between day 21 and day 28. The application of the antiangiogenic drug is at the end of days 21, 23, 25, and 27, and the chemotherapy drug is at the end of days 23, 25, and 27. Our model takes the tumor microenvironment maps just before the initiation of therapy as input and predicts the future tumor microenvironment maps by taking its outputs as new inputs. The predicted tumor microenvironment maps show the effects of ongoing treatment on the tumor microenvironment. A qualitative assessment of Fig 1 shows that our model successfully predicts future tumor density, vasculature, IFP, antiangiogenic drug, and chemotherapy drug maps during ongoing treatment.

**Fig 1.**
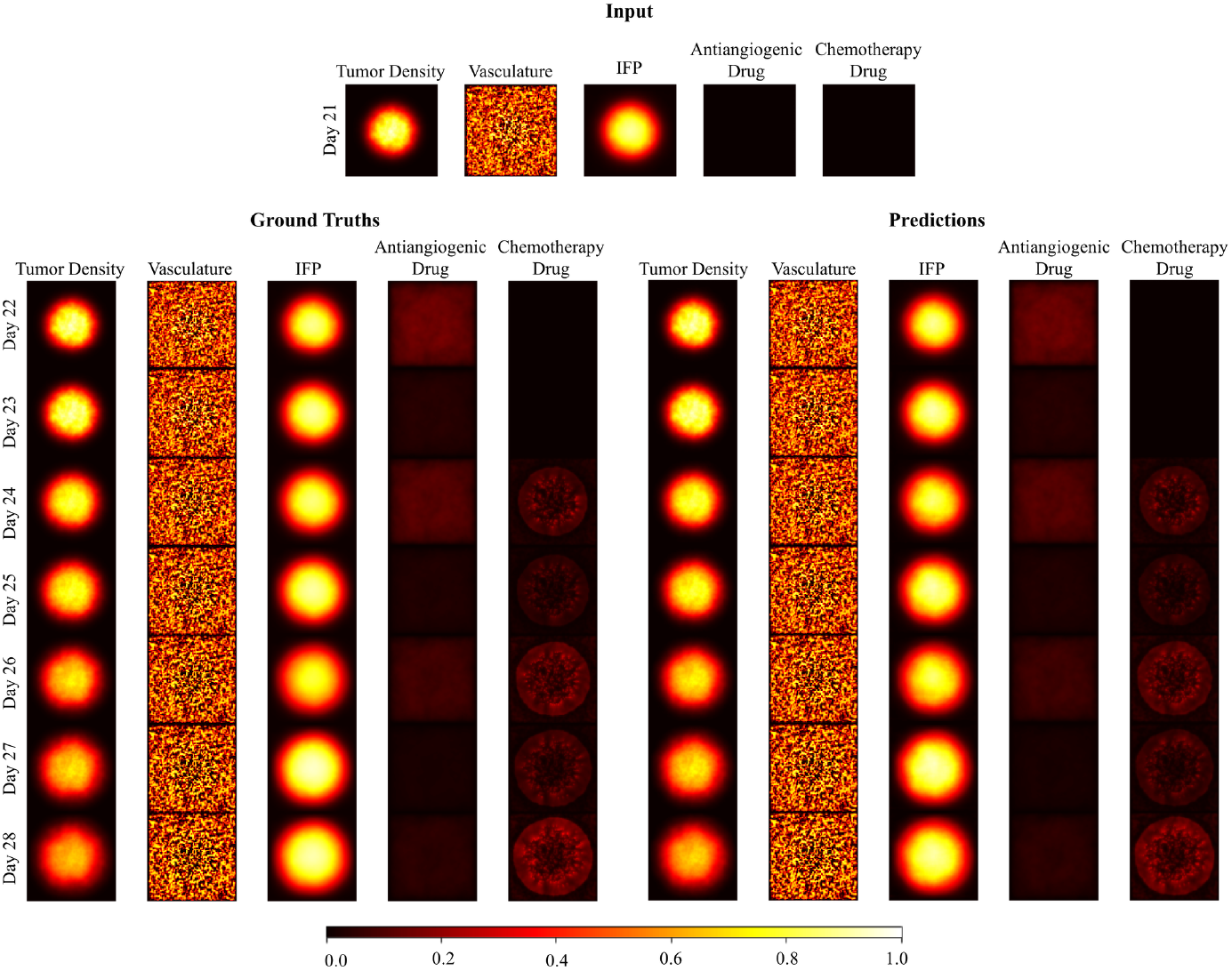
The synthetically generated tumor microenvironment maps of the example patient. The patient receives treatment between days 21 and 28. The application of the antiangiogenic drug is at the end of days 21, 23, 25, and 27 while the chemotherapy drug is applied at the end of days 23, 25, and 27. Drug scalars *A* and *d* have the values of 0.6 and 0.8 respectively. The model takes initial tumor density, vasculature, IFP, antiangiogenic drug, and chemotherapy drug maps as an input (on the top). Ground truth maps that show the ongoing treatment from day 22 to day 28 (on the left). Predicted maps that are iteratively generated from day 22 to day 28 (on the right).

In the experiments, we use 135 test cases from three different patients. We have included five different therapy initiation days and nine different dosage combinations of antiangiogenic and chemotherapy drugs in the cases. Therefore, the cases differ by the treatment initiation days, applied drug dosages, and patient characteristics. A case corresponds to a tensor of 8 days-long tumor microenvironment maps sized 151 × 151 × 8 × 5 (Width x Height x Time Dimension x Channels) accompanied by drug scalars 8 × 2 generated synthetically by using a mathematical model [21]. The consecutive frames correspond to time steps from 0 to 7 and are separated by a day. The five channels represent tumor cell density, vasculature, IFP, antiangiogenic drug diffusion, and chemotherapy drug diffusion maps. Drug scalars denote the dosage for drug insertion in each time step if there is. The insertion schedule is the same for all cases; antiangiogenic drug is inserted at the end of time steps 0, 2, 4, and 6 and chemotherapy drug is inserted at the end of time steps 2, 4 and 6.

By using completely new cases to test our model, we assure that the model does not see the test data before. Each forward step of the DENT model is trained to predict tumor maps at the next time step which is a day ahead. The average PSNR and SSIM scores are shown in Fig 2 over the test set. We see that our model handles to predict future tumor microenvironment maps up to seven days ahead with the average structural similarity score of 0.973 and the average peak signal ratio score of 35.41. The decreasing trends in SSIM and PSNR scores over time are expected as the prediction error is accumulated in each forward step in time. We also observe drops in average SSIM and PSNR scores at time steps 3, 5, and 7 which correspond to one day ahead predictions following the drug injections. Therefore, a possible reason behind these fluctuations may be the challenges in the drug insertion processes.

**Fig 2.**
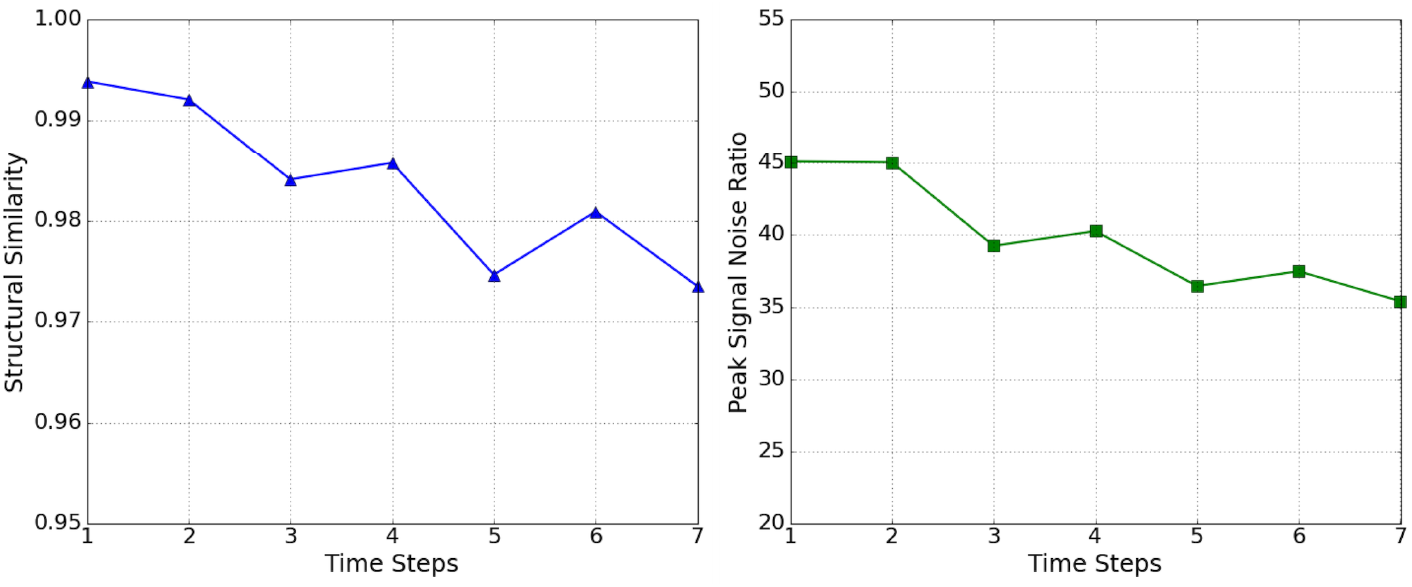
Frame wise average SSIM scores (on the left) and PSNR scores (on the right) over the 135 test cases.

We next examine if the model encapsulates the tumor growth and shrink patterns during the treatment by calculating a dimensionless metric called tumor cell density, *C*. Tumor cell density over the whole tumor shows the effectiveness of ongoing treatment. It is calculated by averaging the pixel values after thresholding the pixels of the tumor density map as performed in the previous study [21]. We compare the tumor cell density of ground truths, *C_gt_*, and predictions, *C_pred_*, over the time steps. For the selected patient case in Fig 1, ground truth and predicted tumor cell density values during treatment are presented in Fig 3. We calculate the absolute percentage error between *C_gt_* and *C_pred_* for 135 test cases which are shown in Table 1. The mean absolute percentage error at the end of day 7 is 2.292 ± 1.820 which shows that our model encapsulates the tumor growth and shrink patterns under the effects of ongoing treatment. It can be seen that the standard deviation of error increases over time since the error accumulates in each prediction.

**Fig 3.**
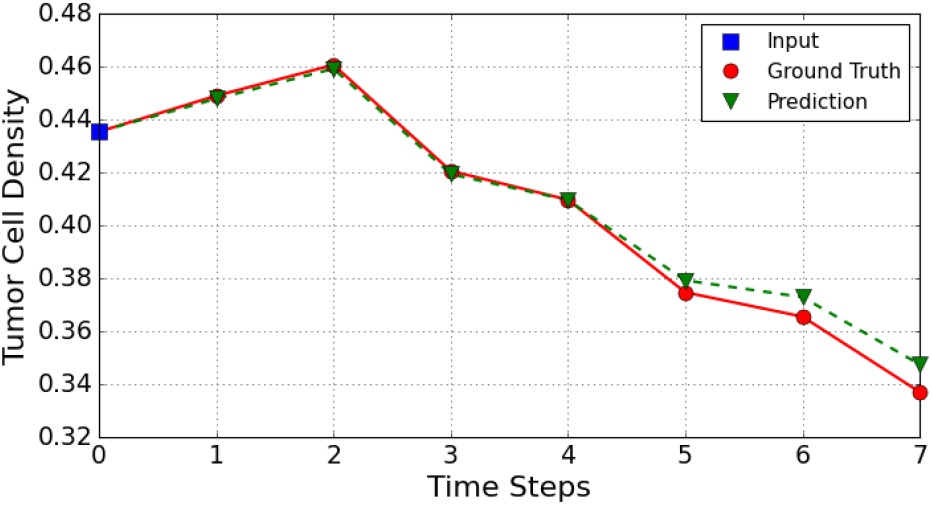
The comparison of tumor cell density values for ground truths and predictions through time steps 0 to 7 for the example case. The tumor cell density at time step 0 corresponds to the tumor cell density of input. (The application of the antiangiogenic drug is at the end of time steps 0, 2, 4, 6 and chemotherapy drug is at the end of time steps 2, 4, 6.)

**Table 1.** Mean absolute percentage error (MAPE) between ground truth and predicted tumor cell densities over days for the test cases.

|  | Day 1 | Day 2 | Day 3 | Day 4 | Day 5 | Day 6 | Day 7 |
| --- | --- | --- | --- | --- | --- | --- | --- |
| MAPE | $0.152 \pm 0.117$ | $0.198 \pm 0.151$ | $0.681 \pm 0.584$ | $1.084 \pm 0.782$ | $1.402 \pm 1.210$ | $2.048 \pm 1.425$ | $2.292 \pm 1.820$ |

## Discussion

In this study, we build a deep learning model designed to solve the chemotherapy response prediction problem. The model is trained with simulation data, which includes tumor growth and angiogenesis, reflecting the biological aspects of tumor microenvironment [20,21]. Here, the proposed deep learning model can predict chemotherapy’s response to tumor structure over an indefinite period via iterating for an indefinite number of times, permissively. Our model operates on a total of five channels: the maps of tumor cell density, vasculature, interstitial fluid pressure, and antiangiogenic and chemotherapy drug diffusions. The predicted maps show the spatial and temporal effects of ongoing treatment on tumor microenvironment maps. The model can take its outputs as inputs, go further in time, and predict the following state of the tumor by iterating this process as many times as required. In these iterations, there is an option of drug injection fed into the model as an input, if it is in the treatment schedule. To include the injection step in the model, first, the drug diffusion maps on the tumor structure are predicted, then these maps are used for the next tumor state prediction in combination with the other inputs such as tumor cell density, vasculature, and interstitial fluid pressure maps. To evaluate the predictive capabilities of our model, we track PSNR and SSIM scores calculated between the ground truth and predicted tumor microenvironment maps during the treatment. Our model can predict future tumor microenvironment maps up to 7 days with the average SSIM score of 0.973 and the average PSNR score of 35.41 at the end of day 7. We have assessed the tumor cell density among the entire predicted tumor density *C_pred_* and compared it with the ground truth cell density *C_gt_.* To obtain the values of cell density both for the predicted tumor and the ground truth tumor, we have used the same method as [21], merely averaging the pixel values that are above a threshold value. We use the percentage error between *C_pred_* and *C_gt_* to evaluate the performance. The model predicts tumor cell density at the end of day 7, with the mean absolute percentage error of 2.292 ±1.820.

Chemotherapy response prediction aims to successfully predict whether the patient will respond to the treatment, depending on the tumor characteristics of the patient. Current approaches to chemotherapy response prediction are mainly based upon classification models. A recent study [48] focuses on the prediction of lung cancer treatment response for patients treated with chemoradiation. By analyzing time-series CT images of patients, the survival probability and cancer-specific outcomes have been predicted. In another study by Ha *et al.* [37], it is aimed to predict the effectiveness of neoadjuvant chemotherapy (NAC) response by using a breast MRI tumor dataset. Patients are classified into three groups as complete, partial, and no response, based on their response to NAC treatment. Although these classification-based models predict chemotherapy response to some extent, their predictions lack spatial and temporal information, which provides the necessary explanations behind predictions. For instance, providing maps of future tumor microenvironment states after the treatment may give clinicians an idea for re-planning the treatment scheduling process. Therefore, there is a challenge in building models that provide detailed information in the chemotherapy response prediction task, which brings us to the modeling of the tumor microenvironment.

On the other hand, mathematical models rely on nonlinear PDEs that simulate the dynamics of the tumor microenvironment, considering the effects of treatments. These models enable us to observe the diffusion of the chemotherapy drug to tumor cells via the normalized vasculature network, and the effectiveness of the applied treatment by producing maps of the future tumor microenvironment states. This approach may eliminate the main drawback of the class-based chemotherapy response prediction models, which is a lack of insight behind the predictions. A recent study [21] presents a mathematical model to simulate the tumor state according to the previous tumor state and the drug diffusion inside the tumor. In each iteration of the model, a predetermined theoretically-thirty-minute gap period proceeds, and the subsequent tumor state is calculated. This allows the model to output the tumor states with a time-resolution of thirty minutes, which is suitable to study the tumor shrinkage or growth comprehensively under the effects of ongoing treatment. The model is deterministic, which means differential equations, predetermined biophysical factors, and constraints are utilized. In potential clinical use, this could hinder the model from making correct interpolations, since these factors and constraints could be varying from patient to patient contrary to the fixed values used in the simulation. Capturing these factors and constraints from patient data is essential to predict patient-specific chemotherapy response. Calculations in the simulation are at the pixel level, and the model cannot capture the detailed physical structure of the tumor and recognize any particular situation, such as detecting the tumor type or distinguishing the smaller sections of the tumor. Since the coefficients used in the differential equations are constant and non-learnable, a deterministic model could not learn from clinical data and be precise enough for clinical use, making it impossible to generalize over thousands of patients.

In the wake of these limitations in the literature, we propose a deep convolutional model. The model simulates tumor microenvironment maps, which encapsulate spatio-temporal effects of the ongoing treatment such as tumor growth and shrinkage patterns, together with the drug diffusion maps. The model outputs future tumor microenvironment maps, which fill the explanatory information gap in classification-based response prediction models. The sequence of predicted maps indicates the response to the therapy in a daily regime. The model predictions reveal the effectiveness of therapy schedule and drug dosages on patients before the application of the therapy. This could allow better adjustment of schedules, use of the potent schedules, and elimination of the ineffective ones. Although mathematical models can simulate tumor microenvironment, they tend to neglect patient-specific tumor response patterns due to a limited number of parameters, fixed coefficients, and deterministic rules governing the model. Since the proposed model is built with end-to-end learning, it can learn patient-specific tumor response patterns in the presence of sufficient data. The model may improve the personalization of treatment by extracting specific clinical features of a patient and making predictions based on them.

Our model for the chemotherapy response prediction task has potential clinical uses; however, some limitations can be eliminated with further improvements. A limitation of the proposed model is using 2D tumor microenvironment maps instead of 3D maps, which can make better use of the tumor microenvironment’s spatial information. When the tumor microenvironment maps are missing the 3rd spatial dimension, the true nature of the tumor cannot be reflected in detail, leading to an incomplete representation of the tumor microenvironment within the model. Therefore, 3D mathematical models of tumor [49] or 3D scanning methods such as CT are the primary candidates of data sources to solve the incomplete environment representation problem. In addition to the 3rd spatial dimension, tumors also have a continuous structure across the time dimension, that is, a continuous course of growth and shrinkage. Modeling the time dependencies between the sequential states of the tumor may be necessary, assuming that the future tumor growth or shrinkage pattern follows its past trend under constant conditions. The input should consist of temporal information of the tumor microenvironment maps to encapsulate the effects of time dependencies on a tumor microenvironment. The model can utilize the input sequence to reveal the features of patient-specific tumor dynamics. These features can be critical to include patient-specific tumor growth patterns and tumor aggressiveness in model predictions. Although our CNN-based model captures spatio-temporal dependencies to some extent, it is preferred to use long short-term memory (LSTM) [50] networks with sequential inputs, combined with CNNs. Convolutional LSTMs [51] can be utilized to extract both spatial and temporal dynamic changes in a single network as they are already exploited in the tumor growth prediction problem [52]. The main challenge of this approach would be to acquire regularly timed and spatially aligned scans to generate sequential data.

The simulated cases used in our study differ by therapy initiation day, drug combinations used in the therapy, and unique patients imposed by the randomly initialized vasculature maps. However, all simulated cases share the same insertion schedule in which antiangiogenic drug therapy is initiated at the beginning of the therapy, and chemotherapy is started with the delay of two days. Both drugs are inserted every other day. Since our cases lack the variations in the insertion schedules, our model may be limited in encapsulating the effects of longer or shorter time intervals between the initiation of antiangiogenic and chemotherapy treatment or non-overlapping insertion days of two types of drugs. A possible solution is incorporating various realistic insertion scheduling scenarios in the cases.

Obtaining the training data from simulations is beneficial in a proof of concept study; however, for a distinguished and genuinely working model, using clinical training data is inevitable. In order to make sure that a deep learning model can simulate and generalize chemotherapy response in a real scenario, it is necessary to train a model with a reasonably large amount of clinical data with sufficient variations of treatment schedules. A clinically applicable response prediction model’s crucial feature is its ability to make predictions in the same domain with clinical data such as MRI, CT, and PET scans. Each scanning technique emphasizes a particular aspect of a biological structure, capturing a unique type of information about the tumor microenvironment. By using these scanning techniques, it is possible to obtain the maps of tumor cells, tumor vasculature [53], IFP [54] and cytotoxic drug distributions [55, 56]. Various scanning techniques should be combined to feed the model’s input to represent the tumor to the full extent, thereby introducing as many aspects of the tumor as possible. This diversity of scan types would provide a complete representation of the tumor within the model, boosting prediction and simulation capabilities.

## Methods

### The mathematical model for acquiring synthetic data

In this study, a mathematical model of tumor built in our previous study [21] is used to obtain the training and testing data sets. Non-linear PDEs are written in dimensionless form and utilized in the model to mimic the tumor and its microenvironment biologically.

The model incorporates tumor cell density and vasculature as well as their interplay and consists of tumor cell density, vasculature, IFP, antiangiogenic agent, and chemotherapy drug. The reaction-diffusion equations are used to describe the spatio-temporal distribution of tumor cell density and vasculature (Eqs 1 and 2). Tumor cell density and vasculature are denoted by n(**x**,t) and m(**x**,t), respectively. In Eq 1, the first term on the right-hand side defines the diffusion of tumor cell density, where *D_n_* is the diffusion coefficient, the second term models the tumor growth rate, where r is the growth rate and *n_l_im* is the maximum carrying capacity. The third term couples the tumor cell density and vasculature in which *α_m_n* indicates the proliferation rate of tumor cells when tumor vessels are present, and the fourth term describes the relationship between the tumor cell density and the chemotherapy drug, here *d_r_* is the elimination rate of tumor cells when chemotherapy drug is applied. Initially, tumor cells are assumed to be distributed by Gaussian.

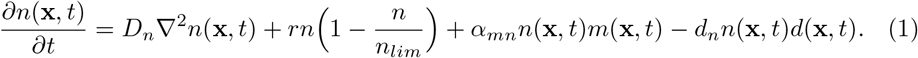

Distinct to healthy vessels, tumor vasculature is a heterogeneous structure, having vessels with tortuous and larger pores. In the model, the Eq (2) is used to represent the heterogeneous tumor vascular network, and it involves the terms for the diffusion of vessels, the production of islands of vessels, the directed motion of vasculature to tumor cells, the production of tumor vessels due to tumor-induced angiogenesis and the elimination of vessels by application of antiangiogenic agent, respectively. For the initial state of tumor vessels, it is assumed that tumor vessels have a random and positively distributed configuration.

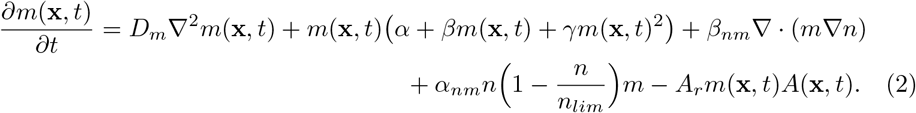

To describe IFP in the solid tumor, the Eq (3) is used:

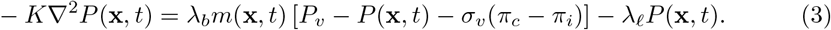

where the term *λ_b_m*(**x**,*t*) [*P_v_— P*(**x**, *t*) — *σ_v_(π_c_—π_i_*)] is the fluid source from blood vessels to the interstitial space and *λ_ℓ_P* (**x**, *t*) is the drainage of fluid from interstitial space to lymph vessels. In the equation, *λ_b_* and *λ_ℓ_* are the hydraulic conductivities of blood and lymp vessels, respectively. The parameter *P_v_* is the vascular pressure, *P* is the interstitial fluid pressure, *σ_v_* is the osmotic reflection coefficient. The terms *π_c_* and *π_i_* indicate the capillary and the interstitial oncotic pressures.

The transport of the antiangiogenic agent is represented by the Eq (4). Here, the first term on the right-hand side is the diffusion of an antiangiogenic agent, where *D_A_* is the diffusion coefficient of the antiangiogenic agent in tissue. The second term is the diffusion of the agent through vessels where *λ_A_* is the transvascular diffusion coefficient of the antiangiogenic agent, and *A_v_* is the concentration of the agent in plasma. The third term is the drainage of agents into the lymphatics, and the last term is the natural decay of agents in tissue, where *k_A_* is the decay rate of antiangiogenic agents.

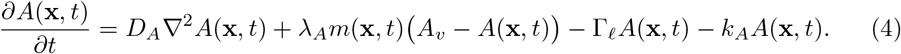

A convection-diffusion equation is used to present the delivery of chemotherapy drug molecules, which are larger particles (around 100 nm in size). In the Eq (5), the first and second terms define the diffusion and convection of chemotherapy drugs in the tissue where *D_d_* is the diffusion coefficient of drugs, and *k*_*E*_ is the retardation coefficient for convection in the interstitium. The third term is the convection of drugs through the vessels, where *σ*_*d*_ is the solvent drag reflection coefficient, the fourth and fifth terms are the drainage of chemotherapy drugs to the lymph vessels and the reaction of chemotherapy drugs with tumor cells, where *d_r_* is the reaction rate. The last term represents the decay of drugs in tissue, where *k_d_* is the natural decay rate.

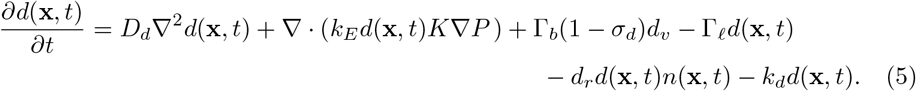

The Eqs (4) and (5) are solved in steady-state since the time scale for the transport of antiangiogenic agent and chemotherapy drug is shorter than the time scale of the tumor growth. Antiangiogenic agent and chemotherapy drug are applied with bolus injection through an exponential decay function:

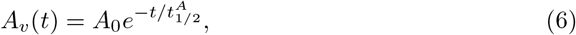

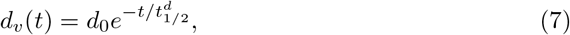

where *A_0_*, *d_0_*, 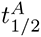 and 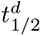 stand for the initial plasma concentration and the plasma half-life of antiangiogenic agent and chemotherapy drug, respectively. No-flux boundary conditions are applied for antiangiogenic agent and chemotherapy drug.

### Datasets

The datasets used in this study generated synthetically by a MATLAB simulation [21] which models tumor microenvironment mathematically as it is described in the previous section. The simulation results are consistent with experimental studies, allowing them to be used for data generation [57–61]. Two datasets are used in the training of the proposed deep learning model DENT. In the training and validation sets, cases from five unique patients are used. A case corresponds to tumor microenvironment maps during eight-day combination therapy treatment. The cases contain tumor microenvironment maps with a time-resolution of 30 minutes. The cases have five different therapy initiation days in the range of day 14 to day 21. The combinations of three different antiangiogenic and chemotherapy drug dosages are used in treatments. The insertion schedule is the same in all treatments. The antiangiogenic drug is inserted at the end of days *t*_0_, *t*_2_, *t*_4_, *t*_6_ and chemotherapy drug is inserted at the end of time steps *t*_2_, *t*_4_, *t*_6_ where *t*_0_ denotes the starting day of therapy. Each patient has 45 cases considering the initiation days and drug dosage combinations. Training and validation datasets are formed by utilizing 225 synthetically generated cases with five unique patients.

The first dataset, employed by Diffuser, contains drug map changes in the presence of drug insertion, a combination of antiangiogenic and chemotherapy drugs (see Fig 4).

**Fig 4.**
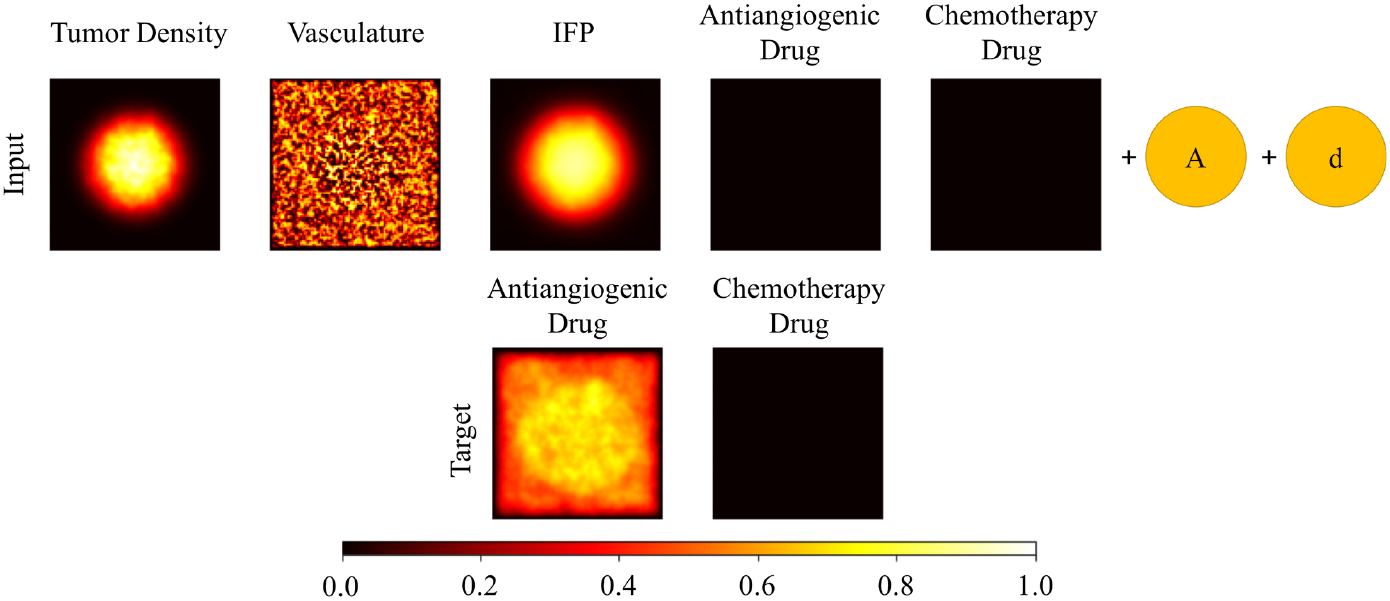
An example pair from the Diffuser dataset. The input and target tensors are presented in first and second rows respectively. The input tensor consists of five channels which are tumor density, vasculature, IFP, antiangiogenic drug, and chemotherapy drug maps accompanied by antiangiogenic drug dosage (A) and chemotherapy drug dosage (d). The target tensor consists of two channels which are antiangiogenic drug and chemotherapy drug maps. The time interval between input and target tensor is an hour.

This dataset consists of 900 pairs. The first element is the tumor microenvironment tensors accompanied by drug scalars that represent the dosage of the given drug. The second element is the drug diffusion maps tensor immediately after drugs are diffused. The size of tumor microenvironment maps in pairs is 151 × 151. It should be noted that we only consider changes in drug diffusion maps between input and target tensors. Therefore, the time interval between the input and output tensors corresponds to an hour in which drugs are fully diffused in the tumor microenvironment. Using these pairs in the Diffuser dataset, we train our model to approximate another mapping function between input and target tensors, which is equivalent to learning dynamics of drug diffusion given the tumor microenvironment maps and drug dosage scalars. Since there is not a considerable change in tumor density, vasculature, and IFP maps in an hour, we omit these channels in the target tensor.

The second dataset, used by Elapser, contains the changes in the tumor microenvironment in the absence of drug insertion between input and target tensors (see Fig 5). The dataset consists of 1575 well-separated pairs of tumor microenvironment maps. The first element of the pair is the input tensor, whereas the second element is the target tensor. Both of the tensors comprise five channels sized by 151 × 151, which are tumor density, vasculature, IFP, antiangiogenic drug, and chemotherapy drug maps. Each pair of tensors is separated by a day apart. By using these pairs, we train our model to approximate a mapping function that maps input tensors to target tensors. Since the time interval between the input and target tensors corresponds to a day, the task is equivalent to learning the dynamics of tumor states in a daily regime.

**Fig 5.**
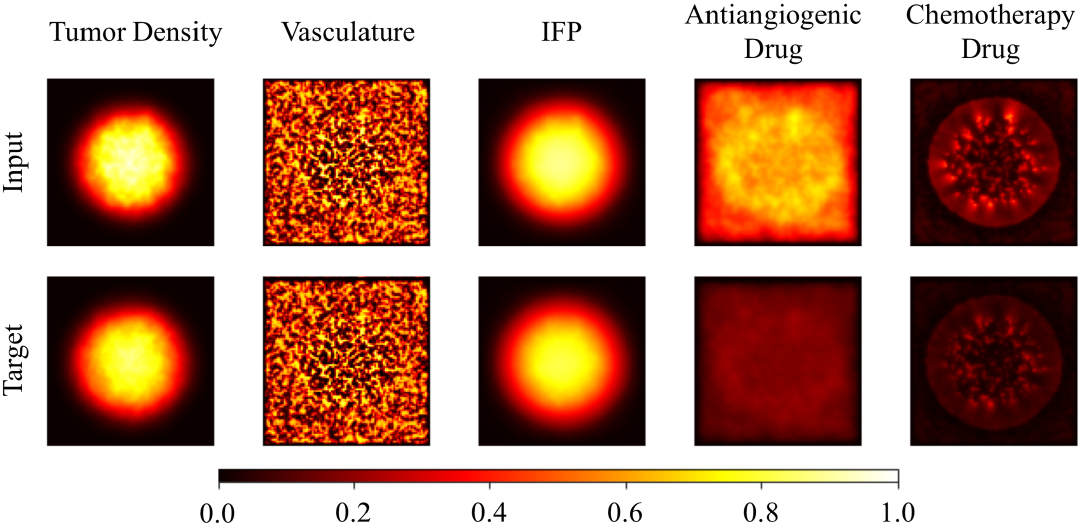
An example pair from the Elapser dataset. The input and target tensors are presented in first and second rows respectively. Each tensor consists of five channels which are tumor density, vasculature, IFP, antiangiogenic drug and chemotherapy drug maps. The time interval between input and target tensor corresponds to a day.

### Data Preprocessing

We aim to eliminate any numerical instabilities among the pixel values of input channels and drug scalars. All input images and scalars in the dataset are scaled with min-max scaling by using the following function:

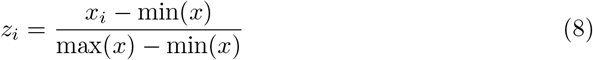

In the case of image inputs, such as tumor density, vasculature, IFP, chemotherapy, antiangiogenic maps, *x_i_* is the pixel value, max(*x*) is the maximum pixel value and min(*x*) is the minimum pixel value in the interesting channel. In the case of scalar inputs, such as antiangiogenic and chemotherapy drug dosages, *x_i_* is drug dosage value, max(*x*) is the maximum value, and min(*x*) is the minimum value from corresponding drug. The normalized value of *x_i_* corresponds to *z_i_*. After the scaling, each pixel and scalar input is within the range of [0,1]. It should be noted that each max(*x*) and min(*x*) values are obtained from the training set to ensure that we do not use any information obtained from the test set.

### The Deep Learning Model

#### Functionality of Simulating an Indefinite Period of Time

The proposed model takes the tumor microenvironment maps *X_t_* and drug scalars *S_t_* and predicts the future microenvironment maps *X*_*t*+1_ in each forward pass. The time difference between *t* and *t* + 1 corresponds to a day. Since *X_t_* and *X*_*t*+1_ are both tumor microenvironment maps, we can feed the model with its prediction *X*_*t*+1_ and *S*_*t*+1_ to generate *X*_*t*+2_. This process can go on indefinitely to obtain the tumor state at any time point *X_t+x_*, where *x* is arbitrary and shows the number of prediction steps.

#### DENT model as the composition of Elapser and Diffuser models

The DENT model has the task of predicting future tumor microenvironment maps considering the effects of ongoing treatment. The model consists of multiple CNN submodels in order to extract spatial features from multichannel tumor microenvironment inputs. The model utilizes these features to predict future tumor microenvironment maps, which is equivalent to learning spatio-temporal dynamics of the tumor microenvironment. The model has two main tasks; the first task is inserting and diffusing the given drug dosages in the given tumor microenvironment, and the second task is predicting future tumor microenvironment maps. Since these tasks are not overlapping, we build two submodels called Diffuser and Elapser networks, which work independently. These two models are integrated to attain the DENT network, which can iterate with taking its outputs as inputs. It is able to check if there is any injection and interfere with the iteration process in case of necessity.

The first part of the proposed model, the Diffuser network, contains two independent CNN submodels, which inserts and diffuses the scalar drug dosages given together with the input tumor microenvironment maps, generating a new drug diffusion maps. It is a conditional network which is initiated if scalars are different from zero. In the case of zero drug dosage, the Diffuser network outputs the relevant drug map without any change. The second part, the Elapser model, contains five independent CNN submodels, which takes tumor microenvironment maps as input tensor and predicts future tumor microenvironment maps in a channel-wise manner. The Elapser model encapsulates the changes in each channel so that it can extrapolate to the future.

By creating a pipeline with the Diffuser and Elapser networks as shown in Fig 6, we build a deep learning model that iterates for a given number of time steps. It takes its outputs as the inputs of the next step while iterating, which is suitable for performing intervention due to drug insertion schedule.

**Fig 6.**
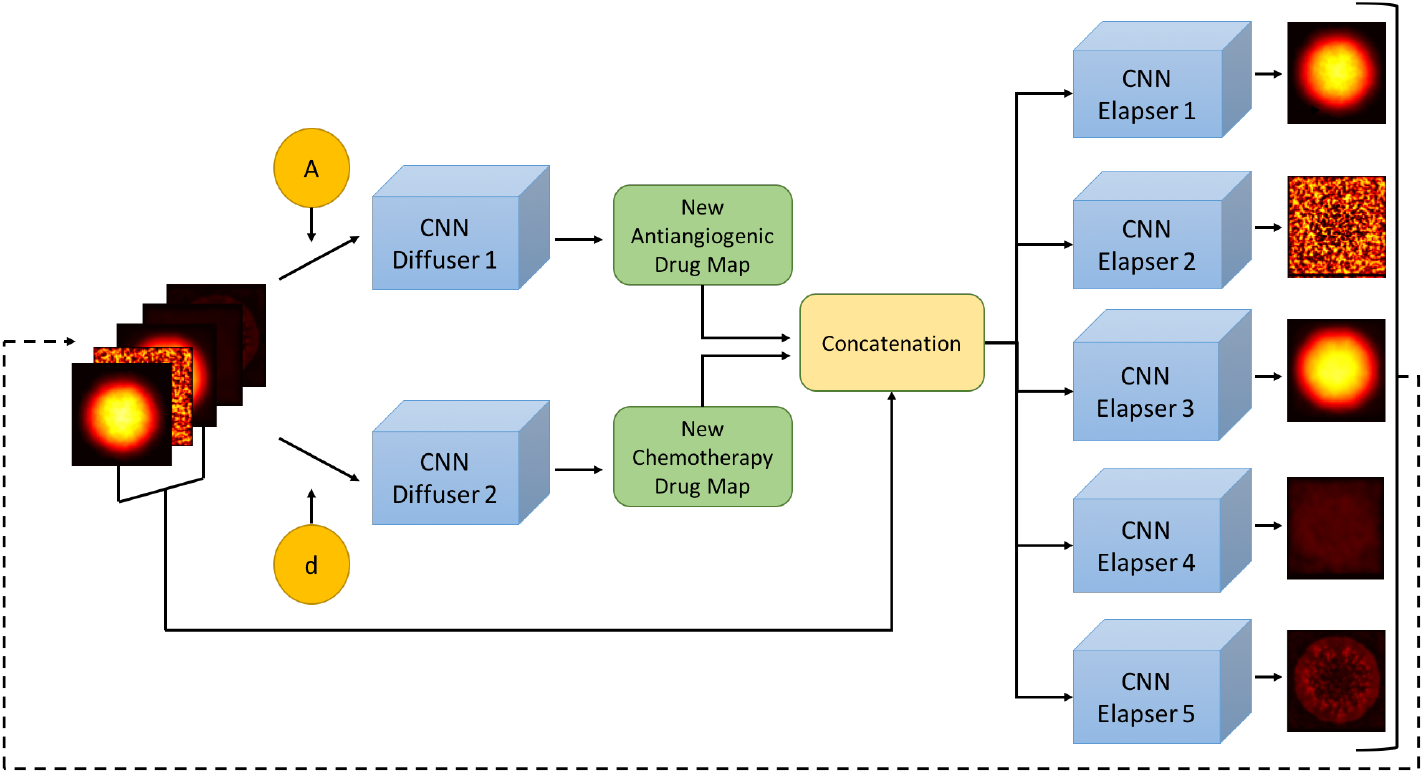
The model diagram. The model takes the five-channel tumor microenvironment maps with antiangiogenic drug dosage (A) and chemotherapy drug dosage (d). The Diffuser submodel predicts new antiangiogenic and chemotherapy drug diffusion maps then the tumor density, vasculature, and IFP maps are concatenated with new drug diffusion maps. The Elapser submodel takes the concatenated tumor microenvironment maps and predicts tumor microenvironment maps of the next day. Since the model has the capacity of using the final prediction as an input, it can predict the tumor microenvironments maps over time.

### Model Specifications and Training Procedures

As previously mentioned, the proposed pipeline consists of two submodels. The first submodel, Diffuser Network, injects and diffuses the scalar drug dosage given the input tensor, generating new antiangiogenic and chemotherapy drug diffusions maps. An input tensor for the Diffuser Network contains tumor microenvironment maps accompanied by two dosage scalars, as shown in Fig 4. We first tile these scalars to our input tensor so that the input has the shape of 7 × 151 × 151. The Diffuser network includes two CNNs that are trained separately with the same input tensors. The first CNN approximates a new antiangiogenic drug diffusion map, and the second CNN approximates a new chemotherapy drug diffusion map. The second submodel, Elapser Network, forecasts the future tumor microenvironment maps by extrapolating from the input tensor shaped 5 × 151 × 151 shown in Fig 5. It outputs the future tumor microenvironment maps as a five-channel image tensor. There is a separate CNN block for predicting each channel of future image tensor.

All CNN submodels are trained separately as their tasks are different from each other. In the training, we use 225 synthetically generated cases from five unique patients. We perform 5-fold cross-validation among the five patients. Therefore, there is not any spatial or temporal dependency between training and validation sets. The submodels are trained with MSE loss for 200 epochs with early-stopping to avoid over-fitting. The activation function is ReLu and kernel size fixed as 5 × 5, whereas stride length is 1. After each layer, 2D batch normalization is performed except the final layer. We use ADAM [45] optimizer with a batch size of 10 and initial learning rate of 10^−4^. The number of layers and filters are selected based on the best performance, and increasing the depth of network or number of filters does not affect the model performance. The model specifications are shown in Table 2. All models are built with Pytorch [46] backend and trained on a single NVIDIA Tesla K80 GPU.

**Table 2.**
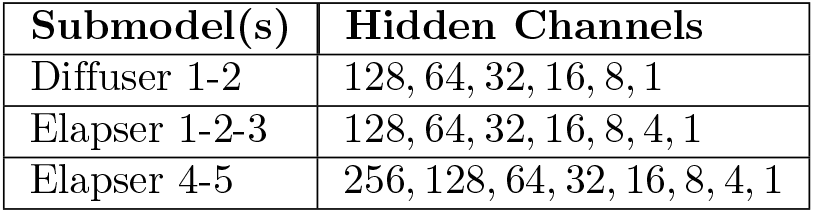
Model Specifications.

| Submodel(s) | Hidden Channels |
| --- | --- |
| Diffuser 1-2 | 128, 64, 32, 16, 8, 1 |
| Elapser 1-2-3 | 128, 64, 32, 16, 8, 4, 1 |
| Elapser 4-5 | 256, 128, 64, 32, 16, 8, 4, 1 |

## References

1. Carmeliet P, Jain RK. Angiogenesis in cancer and other diseases. Nature. 2000;407(6801):249.

2. Jain RK. Normalization of tumor vasculature: an emerging concept in antiangiogenic therapy. Science. 2005;307(5706):58–62.

3. Jain RK. Barriers to drug delivery in solid tumors. Scientific American. 1994;271(1):58–65.

4. Boucher Y, Baxter LT, Jain RK. Interstitial pressure gradients in tissue-isolated and subcutaneous tumors: implications for therapy. Cancer Research. 1990;50(15):4478–4484.

5. Jain RK. Normalizing tumor microenvironment to treat cancer: bench to bedside to biomarkers. Journal of Clinical Oncology. 2013;31(17):2205.

6. Tong RT, Boucher Y, Kozin SV, Winkler F, Hicklin DJ, Jain RK. Vascular normalization by vascular endothelial growth factor receptor 2 blockade induces a pressure gradient across the vasculature and improves drug penetration in tumors. Cancer Research. 2004;64(11):3731–3736.

7. Davis DW, Herbst RS, Abbruzzese JL. Antiangiogenic Cancer Therapy. CRC Press; 2007.

8. Winkler F, Kozin SV, Tong RT, Chae SS, Booth MF, Garkavtsev I, et al. Kinetics of vascular normalization by VEGFR2 blockade governs brain tumor response to radiation: role of oxygenation, angiopoietin-1, and matrix metalloproteinases. Cancer Cell. 2004;6(6):553–563.

9. Hurwitz H, Fehrenbacher L, Novotny W, Cartwright T, Hainsworth J, Heim W, et al. Bevacizumab plus irinotecan, fluorouracil, and leucovorin for metastatic colorectal cancer. New England Journal of Medicine. 2004;350(23):2335–2342.

10. Reck M, von Pawel J, Zatloukal P, Ramlau R, Gorbounova V, Hirsh V, et al. Phase III trial of cisplatin plus gemcitabine with either placebo or bevacizumab as first-line therapy for nonsquamous non-small-cell lung cancer: AVAiL. Journal of Clinical Oncology. 2009;27(8):1227–1234.

11. Segers J, Di Fazio V, Ansiaux R, Martinive P, Feron O, Wallemacq P, et al. Potentiation of cyclophosphamide chemotherapy using the anti-angiogenic drug thalidomide: importance of optimal scheduling to exploit the ‘normalization’ window of the tumor vasculature. Cancer Letters. 2006;244(1):129–135.

12. Huang Y, Stylianopoulos T, Duda DG, Fukumura D, Jain RK. Benefits of vascular normalization are dose and time dependent. Cancer Research. 2013;73(23):7144–7146.

13. Chauhan VP, Stylianopoulos T, Martin JD, Popovic Z, Chen O, Kamoun WS, et al. Normalization of tumour blood vessels improves the delivery of nanomedicines in a size-dependent manner. Nature Nanotechnology. 2012;7(6):383.

14. Wu M, Frieboes HB, Chaplain MA, McDougall SR, Cristini V, Lowengrub JS. The effect of interstitial pressure on therapeutic agent transport: coupling with the tumor blood and lymphatic vascular systems. Journal of Theoretical Biology. 2014;355:194–207.

15. McDougall SR, Anderson A, Chaplain M, Sherratt J. Mathematical modelling of flow through vascular networks: implications for tumour-induced angiogenesis and chemotherapy strategies. Bulletin of Mathematical Biology. 2002;64(4):673–702.

16. McDougall SR, Anderson AR, Chaplain MA. Mathematical modelling of dynamic adaptive tumour-induced angiogenesis: clinical implications and therapeutic targeting strategies. Journal of Theoretical Biology. 2006;241(3):564–589.

17. Welter M, Rieger H. Interstitial fluid flow and drug delivery in vascularized tumors: a computational model. Plos One. 2013;8(8):e70395.

18. Stephanou A, McDougall SR, Anderson AR, Chaplain MA. Mathematical modelling of flow in 2D and 3D vascular networks: applications to anti-angiogenic and chemotherapeutic drug strategies. Mathematical and Computer Modelling. 2005;41(10):1137–1156.

19. Wu J, Long Q, Xu S, Padhani AR. Study of tumor blood perfusion and its variation due to vascular normalization by anti-angiogenic therapy based on 3D angiogenic microvasculature. Journal of Biomechanics. 2009;42(6):712–721.

20. Kohandel M, Kardar M, Milosevic M, Sivaloganathan S. Dynamics of tumor growth and combination of anti-angiogenic and cytotoxic therapies. Physics in Medicine & Biology. 2007;52(13):3665.

21. Yonucu S, Yılmaz D, Phipps C, Unlu MB, Kohandel M. Quantifying the effects of antiangiogenic and chemotherapy drug combinations on drug delivery and treatment efficacy. Plos Computational Biology. 2017;13(9):e1005724.

22. Eo JS, Jeong JM. Angiogenesis imaging using 68Ga-RGD PET/CT: therapeutic implications. In: Seminars in Nuclear Medicine. vol. 46. Elsevier; 2016. p. 419–427.

23. Barrett T, Kobayashi H, Brechbiel M, Choyke PL. Macromolecular MRI contrast agents for imaging tumor angiogenesis. European Journal of Radiology. 2006;60(3):353–366.

24. Zhang HF, Maslov K, Stoica G, Wang LV. Functional photoacoustic microscopy for high-resolution and noninvasive in vivo imaging. Nature Biotechnology. 2006;24(7):848–851.

25. Hu S, Wang LV. Photoacoustic imaging and characterization of the microvasculature. Journal of Biomedical Optics. 2010;15(1):011101.

26. Horiguchi A, Shinchi M, Nakamura A, Wada T, Ito K, Asano T, et al. Pilot study of prostate cancer angiogenesis imaging using a photoacoustic imaging system. Urology. 2017;108:212–219.

27. Lao Y, Xing D, Yang S, Xiang L. Noninvasive photoacoustic imaging of the developing vasculature during early tumor growth. Physics in Medicine & Biology. 2008;53(15):4203.

28. Hu S, Oladipupo S, Yao J, Santeford AC, Maslov K, Kovalski J, et al. Optical-resolution photoacoustic microscopy of angiogenesis in a transgenic mouse model. In: Photons Plus Ultrasound: Imaging and Sensing 2010. vol. 7564; 2010. p. 756406.

29. Swanson KR, Alvord Jr EC, Murray J. A quantitative model for differential motility of gliomas in grey and white matter. Cell Proliferation. 2000;33(5):317–329.

30. Hogea C, Davatzikos C, Biros G. An image-driven parameter estimation problem for a reaction-diffusion glioma growth model with mass effects. Journal of Mathematical Biology. 2008;56(6):793–825.

31. Roque T, Risser L, Kersemans V, Smart S, Allen D, Kinchesh P, et al. A dce-mri driven 3-d reaction-diffusion model of solid tumor growth. IEEE Transactions on Med Imaging. 2017;37(3):724–732.

32. Sahiner B, Chan HP, Petrick N, Wei D, Helvie MA, Adler DD, et al. Classification of mass and normal breast tissue: a convolution neural network classifier with spatial domain and texture images. IEEE Transactions on Med Imaging. 1996;15(5):598–610.

33. Kallenberg M, Petersen K, Nielsen M, Ng AY, Diao P, Igel C, et al. Unsupervised deep learning applied to breast density segmentation and mammographic risk scoring. IEEE Transactions on Medical Imaging. 2016;35(5):1322–1331.

34. Shin HC, Roth HR, Gao M, Lu L, Xu Z, Nogues I, et al. Deep convolutional neural networks for computer-aided detection: CNN architectures, dataset characteristics and transfer learning. IEEE Transactions on Medical Imaging. 2016;35(5):1285–1298.

35. Zhang L, Lu L, Summers RM, Kebebew E, Yao J. Convolutional Invasion and Expansion Networks for Tumor Growth Prediction. IEEE Transactions on Medical Imaging. 2018;37(2):638–648.

36. Urban G, Bache K, Phan DTT, Sobrino A, Shmakov AK, Hachey SJ, et al. Deep Learning for Drug Discovery and Cancer Research: Automated Analysis of Vascularization Images. IEEE/ACM Transactions on Computational Biology and Bioinformatics. 2019;16(3):1029–1035.

37. Ha R, Chin C, Karcich J, Liu MZ, Chang P, Mutasa S, et al. Prior to initiation of chemotherapy, can we predict breast tumor response? Deep learning convolutional neural networks approach using a breast MRI tumor dataset. Journal of Digital Imaging. 2019;32(5):693–701.

38. Ypsilantis PP, Siddique M, Sohn HM, Davies A, Cook G, Goh V, et al. Predicting response to neoadjuvant chemotherapy with PET imaging using convolutional neural networks. Plos One. 2015;10(9):e0137036.

39. Cha KH, Hadjiiski L, Chan HP, Weizer AZ, Alva A, Cohan RH, et al. Bladder Cancer Treatment Response Assessment in CT using Radiomics with Deep-Learning. Scientific Reports. 2017;7(1):8738. doi:https://doi.org/10.1038/s41598-017-09315-w.

40. LeCun Y, Bengio Y, Hinton G. Deep learning. Nature. 2015;521(7553):436–444.

41. Goodfellow I, Bengio Y, Courville A. In: Deep Learning. MIT Press; 2016.

42. Krizhevsky A, Sutskever I, Hinton GE. ImageNet Classification with Deep Convolutional Neural Networks. In: Advances in Neural Information Processing Systems. vol. 25. Curran Associates, Inc.; 2012. p. 1097–1105.

43. Karpathy A, Toderici G, Shetty S, Leung T, Sukthankar R, Fei-Fei L. Large-scale video classification with convolutional neural networks. In: Proceedings of the IEEE Conference on Computer Vision and Pattern Recognition; 2014. p. 1725–1732.

44. Razzak MI, Naz S, Zaib A. Deep learning for medical image processing: Overview, challenges and the future. In: Classification in BioApps. Springer; 2018. p. 323–350.

45. Kingma DP, Ba J. Adam: A Method for Stochastic Optimization; 2014. Preprint at http://arxiv.org/abs/1412.6980.

46. Paszke A, Gross S, Massa F, Lerer A, Bradbury J, Chanan G, et al. PyTorch: An Imperative Style, High-Performance Deep Learning Library. In: Advances in Neural Information Processing Systems. vol. 32. Curran Associates, Inc.; 2019. p. 8026–8037.

47. Wang Z, Bovik AC, Sheikh HR, Simoncelli EP. Image quality assessment: from error visibility to structural similarity. IEEE Transactions on Image Processing. 2004;13(4):600–612.

48. Xu Y, Hosny A, Zeleznik R, Parmar C, Coroller T, Franco I, et al. Deep Learning Predicts Lung Cancer Treatment Response from Serial Medical Imaging. Clinical Cancer Research. 2019;25(11):3266–3275. doi:https://doi.org/10.1158/1078-0432.ccr-18-2495.

49. Tang L, van de Ven AL, Guo D, Andasari V, Cristini V, Li KC, et al. Computational Modeling of 3D Tumor Growth and Angiogenesis for Chemotherapy Evaluation. Plos One. 2014;9(1):1–12. doi:https://doi.org/10.1371/journal.pone.0083962.

50. Hochreiter S, Schmidhuber J. Long Short-Term Memory. Neural Computation. 1997;9(8):1735–1780.

51. Shi X, Zhourong C, Hao W, Dit-Yan Y, Wai-Kin W, Wang-chun W. Convolutional LSTM Network: A Machine Learning Approach for Precipitation Nowcasting. In: Advances in Neural Information Processing Systems. vol. 28. Curran Associates, Inc.; 2015. p. 802–810.

52. Zhang L, Lu L, Wang X, Zhu RM, Bagheri M, Summers RM, et al. Spatio-Temporal Convolutional LSTMs for Tumor Growth Prediction by Learning 4D Longitudinal Patient Data. IEEE Transactions on Medical Imaging. 2020;39(4):1114–1126.

53. Jackson A, O’Connor JP, Parker GJ, Jayson GC. Imaging tumor vascular heterogeneity and angiogenesis using dynamic contrast-enhanced magnetic resonance imaging. Clinical Cancer Research. 2007;13(12):3449–3459.

54. Hassid Y, Furman-Haran E, Margalit R, Eilam R, Degani H. Noninvasive magnetic resonance imaging of transport and interstitial fluid pressure in ectopic human lung tumors. Cancer Research. 2006;66(8):4159–4166.

55. Saito R, Bringas JR, McKnight TR, Wendland MF, Mamot C, Drummond DC, et al. Distribution of liposomes into brain and rat brain tumor models by convection-enhanced delivery monitored with magnetic resonance imaging. Cancer Research. 2004;64(7):2572–2579.

56. Mardor Y, Rahav O, Zauberman Y, Lidar Z, Ocherashvilli A, Daniels D, et al. Convection-enhanced drug delivery: increased efficacy and magnetic resonance image monitoring. Cancer Research. 2005;65(15):6858–6863.

57. Stapleton S, Milosevic M, Tannock IF, Allen C, Jaffray DA. The intra-tumoral relationship between microcirculation, interstitial fluid pressure and liposome accumulation. Journal of Controlled Release. 2015;211:163–170.

58. Tailor TD, Hanna G, Yarmolenko PS, Dreher MR, Betof AS, Nixon AB, et al. Effect of pazopanib on tumor microenvironment and liposome delivery. Molecular Cancer Therapeutics. 2010;9(6):1798–1808.

59. Graff B, Kvinnsland Y, Skretting A, Rofstad EK. Intratumour heterogeneity in the uptake of macromolecular therapeutic agents in human melanoma xenografts. British Journal of Cancer. 2003;88(2):291–297.

60. Seynhaeve AL, Hoving S, Schipper D, Vermeulen CE, Aan de Wiel-Ambagtsheer G, van Tiel ST, et al. Tumor necrosis factor *α* mediates homogeneous distribution of liposomes in murine melanoma that contributes to a better tumor response. Cancer Research. 2007;67(19):9455–9462.

61. Cabral H, Matsumoto Y, Mizuno K, Chen Q, Murakami M, Kimura M, et al. Accumulation of sub-100 nm polymeric micelles in poorly permeable tumours depends on size. Nature Nanotechnology. 2011;6(12):815.

